# Exposure to aldehyde cherry e-liquid flavouring and its vape by-product disrupts pulmonary surfactant biophysical function

**DOI:** 10.1101/2023.09.22.558976

**Authors:** Alexia Martin, Carmelo Tempra, Yuefan Yu, Juho Liekkinen, Roma Thakker, Hayoung Lee, Berta de Santos Moreno, Ilpo Vattulainen, Christos Rossios, Matti Javanainen, Jorge Bernardino de la Serna

**Affiliations:** National Heart and Lung Institute, Imperial College London, Sir Alexander Fleming Building, London SW7 2AZ, U.K.; Institute of Organic Chemistry and Biochemistry, Czech Academy of Sciences, CZ-16000 Praha 6, Czech Republic; Department of Physics, University of Helsinki, FI-00560 Helsinki, Finland; Institute of Biotechnology, University of Helsinki, FI-00790 Helsinki, Finland

## Abstract

Over the last decade there has been a surge in vaping device usage, especially among adolescents, raising concerns for potentially related lung damage. Notoriously, there have been many e-cigarette or vaping-related lung injury (EVALI) cases resulting in hospitalisations and deaths. Although the vaping component vitamin E acetate has been linked to a large proportion of EVALI cases resulting in its widespread banning, one fifth of the cases remain unexplained. Furthermore, the overall long-term impact of vaping on respiratory health is poorly understood. A likely driver behind EVALI is pulmonary surfactant disruption, as it is the first point of contact for any inhaled toxicant in the alveoli, and abnormalities of its function are linked to some symptoms presented in EVALI cases. Aberrant biophysical function of the surfactant results in alveolar surface tension increase, causing alveolar collapse. Vaping chemicals with the potential to disrupt surfactant function must be hydrophobic molecules able to interact with surfactant components at the alveolar air–liquid interface. Recent findings have recorded the synthesis of highly hydrophobic acetal by-products of the base vaping chemical propylene glycol and common flavouring aldehydes, including the cherry flavouring benzaldehyde, not identified in preliminary e-liquid safety tests. This study provides evidence that benzaldehyde and its by-product, benzaldehyde propylene glycol acetal, have the potential to significantly disrupt surfactant biophysical function *via* interactions with surfactant proteins SP-B and/or SP-C, which may provide stable interactions within the surfactant film by forming associations with the sublayer of surfactant three-dimensional structure present at high lateral compression, *i.e.*, expiration breathing. Data also suggest considerable vaping chemical loss to the experimental subphase, indicating potential further implications to the alveolar epithelial layer beneath.

## Introduction

Ever since their introduction to the market in 2006,^1^ vaping devices have been rapidly growing in popularity. This is largely due to their perceived safety and the addition of flavourings extending the intended target market from adult cigarette smokers to include adolescents.^2^ This strategy has turned out to be treacherously efficient as 27.5% of US high schoolers in 2019 admitted to having vaped in the last 30 days.^3,4^ E-cigarettes are widely accepted to be a safer alternative to smoking,^4,5^ in particular causing less adverse effects on non-lung organs compared to traditional cigarettes.^4^ Still, alongside the surge in their use in the US and UK has come a rise in vaping-related lung injuries resulting in hospitalisations and deaths, ^4,6,7^ with 2807 total hospitalised cases recorded by February 2020.^1^ Alarmingly, 78% of those admitted were under 35 years old, showcasing the danger facing the younger population.^8^ These patients were diagnosed with e-cigarette or vaping related lung injury (EVALI), which is assigned when a patient presents with pulmonary infiltrate, is clear of any microbial infection, and has vaped within 90 days.^9^ Pathologically, this corresponds to diffuse alveolar damage, fibrinous pneumonitis, and pneumonia,^10^ with treatment limited to reducing the inflammation *via* corticosteroid administration.^11^ Most cases have been linked to a dilutant used in tetrahydrocannabinol e-liquids; vitamin E acetate (VEA).^6^ This hydrophobic molecule was found to accumulate in the alveoli to cause lipoid pneumonia^12^ and to disrupt the pulmonary surfactant,^13–15^ hence resulting in widespread VEA bans.^16^ Despite this discovery, 20% of EVALI cases remain unexplained,^17^ and the long-term effects of many vaping components are still poorly understood.^4,18^

E-cigarette vapour is known to reach the alveoli, where any inhaled toxicants must first pass the delicate pulmonary surfactant film that sits atop the alveolar liquid. Surfactant has the vital biophysical function of reducing surface tension of the air–liquid interface in the alveoli.^19,20^ Without functioning surfactant, high surface tension would prevent reexpansion after alveolar compression, resulting in alveolar collapse.^21,22^ Aberrant surfactant function and severe alveolar collapse are associated with Acute Respiratory Distress Syndrome (ARDS) which has been the final diagnosis of many advanced EVALI cases ^6,15^ due to shared symptoms of alveolar damage and inflammation.^23^

Pulmonary surfactant is a membranous lipoprotein film synthesised by alveolar type II cells, which forms a monolayer with associated bilayers beneath in the aqueous subphase.^24^ It contains a mixture of lipids (90% of total mass) and proteins (10% of total mass). On the lipid side, 1,2-dipalmitoyl-*sn*-glycero-3-phosphocholine (DPPC) is the most essential component for the reduction of surface tension, as it is the only lipid able to reach a compact gel-like liquid condensed state at physiological temperatures.^25,26^ Phospholipids with unsaturated acyl chains, such as 1-palmitoyl-2-oleoyl-*sn*-glycero-3-phospho-choline and -glycerol (POPC and POPG, respectively) prevent lipid fragility at high compression level by providing sufficient fluidity to the high content of saturated lipids together with cholesterol.^27,28^ For additional film stabilisation, the hydrophobic surfactant proteins SP-B and SP-C—alongside with their roles in gas exchange^29^—work alongside unsaturated lipids to prevent film collapse^30,31^ and thereby material loss. This is achieved by allowing the two-dimensional (2D) surfactant monolayer to fold into a three-dimensional (3D) structure at high compression levels through inter-layer cross-links.^32,33^ Lipids with unsaturated acyl chains and proteins associate with the 3D sublayer reservoir while phospholipids with saturated acyl chains— mainly DPPC—remain at the interface to self-assemble into tight lateral compaction yielding near-gel structure, permitting the attainment of low surface tensions.^24,34,35^ The remaining hydrophilic surfactant proteins SP-A and SP-D are not surface-active, rather they play a role in the innate immune response. ^36^

So far, studies on the effect of vaping on the biophysical properties of pulmonary surfactant have had limited focus on specific components and the conclusions have been somewhat inconsistent; some studies reported minimal to no surfactant disruption,^15,37,38^ whereas others reported the opposite.^39,40^ Therefore, the biophysical impact of flavouring chemicals remains predominantly unexplored, which is concerning as this is considered the main attraction for the younger demographic.^2,18^

Common vaping flavourings include aldehydes, such as the cherry flavouring benzaldehyde (BA), which are widely used in the food and cosmetics industries.^41^ However, their longterm impact on health in a vaping context has not yet been thoroughly explored.^4^ Recent studies found that vaping liquids containing aldehyde flavourings are chemically unstable, ^42,43^ resulting in harmful by-products not recorded in initial e-liquid product safety screenings.^5,18^ It was recently reported that during both storage and the heating process of vaporisation, the base component propylene glycol (PG) and flavouring aldehydes react to form the respiratory irritant PG acetals, for example, benzaldehyde PG acetal (BPGA). It was observed that 40% of BA was converted to BPGA, with a carry-over rate of 50–80%.^42^ Considering BPGA is a highly hydrophobic molecule, ^44^ it was hypothesised that BPGA would sit at the air– liquid interface and interact with surfactant molecules to disrupt biophysical function. The effects of BA on surfactant are also unknown, hence it is being tested alongside BPGA while serving as an additional control as a smaller and less hydrophobic molecule,^44^ and thereby theoretically inducing a weaker effect.

Therefore, the aim of this study was to understand if and how the flavouring aldehyde BA and its by-product BPGA disrupt the biophysical function of the pulmonary surfactant. To this end, the surface activity and their interactions with surfactant constituents were investigated for BA and BPGA, employing both a protein-free synthetic lipid surfactant (SLS)^32,33^ and a clinical surfactant containing also surfactant proteins SP-B and SP-C (Alveofact).^45^ The SLS was developed after lipidomic analysis of human and murine-derived surfactant^27,46^ and served as a control for vaping chemical–protein interactions. For this purpose, two different dynamic compression–expansion surfactometer models were utilised to investigate biophysical function: a quasi-static model (*i.e.*, Langmuir–Blodgett trough compression or cycles, LBT)^47^ and a model replicating physiological dynamics (*i.e.*, constrained sessile drop, CSD).^48^ We monitored changes in the biophysical function of the surfactant formulations by quantifying three parameters, namely the minimum surface tension, compressibility modulus, and hysteresis. To prevent alveolar collapse, a well-functioning human pulmonary surfactant brings the surface tension to below 2 mN/m, has a compressibility modulus that enables high compaction without film collapse, and demonstrates low hysteresis through compression–expansion cycles, defined as limited material loss and efficient lipid re-organisation.^28,49,50^ Alterations in these three functional properties were observed in the presence of vaping chemicals, with a noteworthy implication of the hydrophobic surfactant proteins. Molecular level insight into the interactions of BA and BPGA with surfactant lipids and proteins was obtained from atomistic molecular dynamics simulations. They revealed that both BA and BPGA partitioned to the surfactant monolayer and perturbed its structure. In simulations containing the surfactant proteins, the vaping chemicals accumulated in the vicinit of the hydrophobic proteins in the compressed monolayer, which explains their significant role observed in experiments. Overall, we provide novel molecular insights about how a flavouring aldehyde inhaled from vaping chemicals and its *de novo* by-product impacts surfactant function, which explains plausible toxicological mechanisms that would have implications on respiratory health.

## Results

### BA and BPGA Interfere with Lipids to Compromise the Biophysical Function of the Surfactant

It is known that changes in the lipid composition or variations from the evolutionary refined ratio of saturated and unsaturated chains of lipids and cholesterol drastically influence the biophysical function of lung surfactant monolayers,^27,30^ multilayered films, and the dynamics and inter-connections between the two.^24,31^ To determine the molecular interactions between components of the SLS—which consists of the major surfactant lipids—and the vaping components BA and BPGA, we quantified its biophysical function by means of monitoring the surface pressure at the air–liquid interface on a Langmuir–Blodgett trough (LBT).

We performed 10 consecutive compression–expansion cycles (Π *− A* iso-cycles), as monolayer constituent refinement is known to be an important property of a lung surfactant^20^ **(Supplementary Fig. S1)**. Quasi-static LBT compression rates allow the observation of fine molecular interactions.^51,52^ In the Π *− A* isotherm measured during the first cycle, there was a clear decrease in the Π_max_ when either BA or BPGA was added (Fig. 1A). Therefore, the interaction of BA and BPGA with surfactant lipids prevents the achievement of higher surface pressures. Another interesting aspect to note was that below 10 mN/m—where a liquid expanded (L_e_)-like phase could be expected—neither BA nor BPGA seemed to perturb the surface pressure. This is evident from the similar slopes of increase in the surface pressure measured in the absence or presence of either BA or BPGA. However, once the liquid condensed (L_c_)-like phase is reached at surface pressures *≈*50 mN/m or above—where possible multilayered material could be associated with the interfacial monolayer—vaping components diminished the effect as can be observed by a smoothing of the kinks between 48 mN/m and 63 mN/m. Notably, this effect was less prominent in the last cycle isotherms (Fig. 1B). After a cycling process and refinement of the monolayer, the Π_max_ was significantly reduced and the kinks also vanished (Fig. 1B,C).

**Figure 1:**
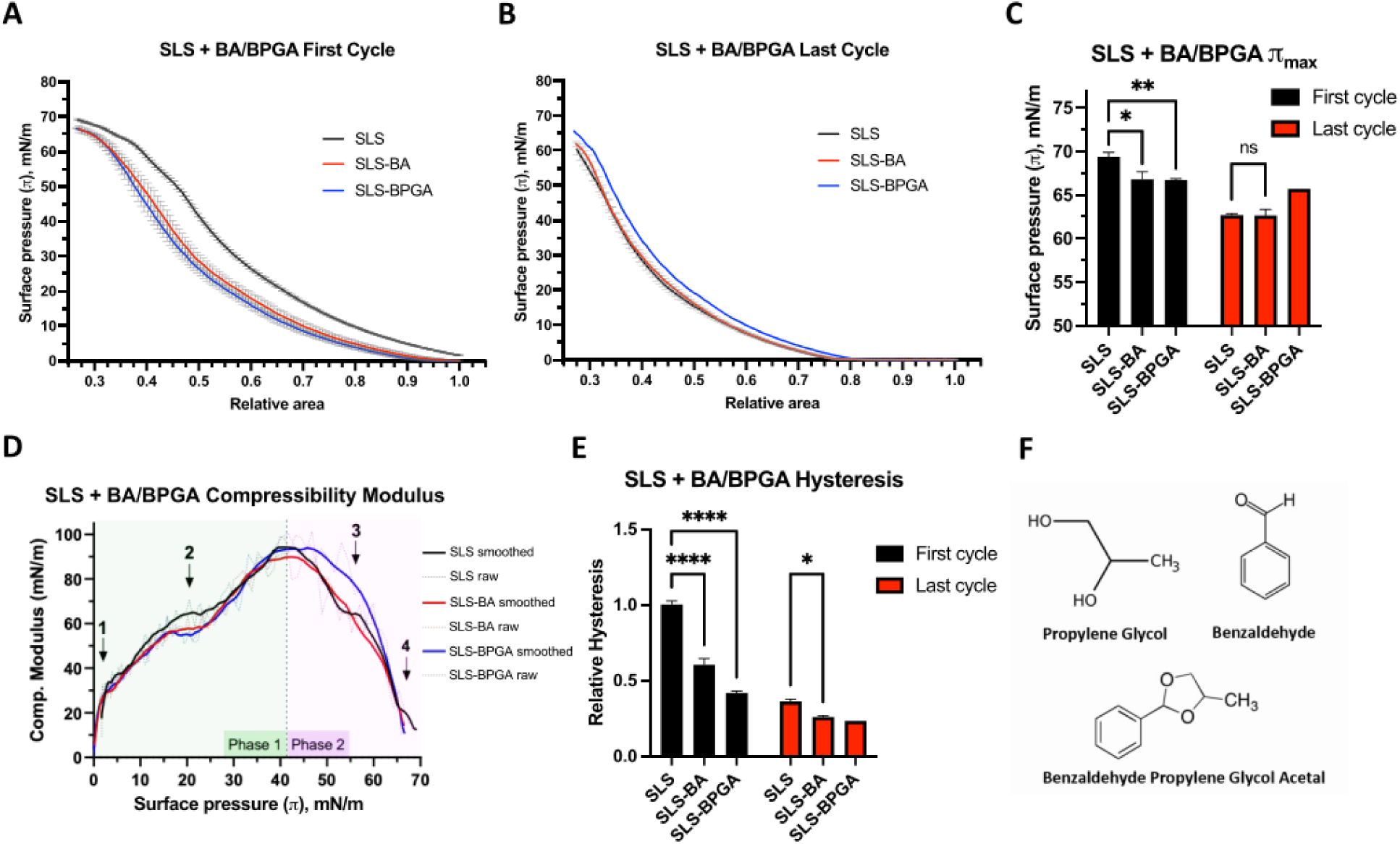
Molecular interaction study between SLS and BA or BPGA. Results from 10 quasi-static compression–expansion LBT iso-cycles. A) LBT Π *− A* isotherms from the first compression–expansion isocycle, N=3. B) Last cycle isotherm (out of a total of 10). SLS and SLS–BA; N=3. SLS–BPGA; N=1. C) Comparison of maximum surface pressures at the first and last (10^th^) cycles. D) Compressibility modulus with both both raw and smoothed data (Savitzky–Golay filter over 9-points) shown. Arrows point to areas of interest on the graph (see text). E) Hysteresis of the first and last cycles, relative to SLS first cycle. Experiments done at 25*^◦^*C. Preliminary results at 37*^◦^*C **(Supplementary Fig. S2)**, did not allow to investigate molecular interactions at high surface pressures. F) Investigated vaping compounds. Significance values represent results from Two-Way ANOVAs; “ns”: not significant (p*>*0.05), *p*<*0.05, **p*<*0.01, ****p*<*0.0001.

Ideally, the compressibility modulus (*κ*) is high leading up to maximum surface pressures, as tight lateral compaction is essential to reach Π_max_. Conversely, lower *κ* is favourable at high surface pressures, as film elasticity prevents collapse—and thereby material loss—by better enabling 2D to 3D transitions. In a protein-free model, mainly unsaturated lipids would move into the 3D reservoir at high surface pressures, especially beyond the “squeeze-out” plateau.^53^ The compressibility modulus was calculated throughout the first cycle exclusively, as this is where it is best defined. ^49,54^ We observed a biphasic behaviour with alterations to SLS compressibility throughout both phases seen with BA and BPGA addition (Fig. 1D). During the incline towards maximum compressibility modulus (Phase 1 in Fig. 1D), *κ* was smaller at smaller surface pressures with BA or BPGA present as compared with SLS alone (point 1 in Fig. 1D). At a surface pressure of 20 mN/m (point 2), BA and BPGA reduce SLS *κ* from 70 mN/m to 58 mN/m and 56 mN/m, respectively. In the decline (Phase 2), SLS–BPGA displays a slower decrease in compressibility modulus than SLS and SLS–BA, increasing *κ* by over 10 mN/m at Π = 55 mN/m (point 3). At a surface pressure larger than 65 mN/m (point 4), both vaping chemicals present loss of the characteristic SLS “kink”, depicting a solid gel phase of lipid compaction which enables SLS to reach 70 mN/m Π_max_. Overall, BA and BPGA negatively influence surfactant lipid compressibility modulus *κ* at high and low surface pressures.

To sustain high surface pressures over multiple cycles, the loss of surfactant constituents from the air–liquid interface at high surface pressures must be avoided, while allowing monolayer refinement to optimise film organisation.^28^ Low hysteresis reflects minimal loss of surface-active material between compression and expansion. That said, BA and BPGA both significantly decrease first cycle SLS hysteresis (Fig. 1E). We observed a 40% and 60% decrease with BA and BPGA, respectively, with both p-values below 0.0001. After ten isocycles, hysteresis had decreased by over 60% in all groups, indicating refinement. Notably, variation between groups was also drastically reduced, with the SLS–BA p-value increasing to above 0.01. As such, BA and BPGA interfere with initial surfactant lipid hysteresis. In summary, the three biophysical function parameters confirm fine interactions between surfactant lipids and BA and BPGA molecules, however these seemed unstable over multiple quasi-static iso-cycles.

### BPGA Partitions to the Acyl Chain Region of SLS Monolayers and Increases their Packing

Having observed that BA and BPGA interactions with surfactant lipids at continuous quasistatic compression detrimentally influence the functional surface pressure of SLS monolayers, we set on to resolve the possible molecular mechanisms that would lead to this behaviour. An obvious element of differential behaviour between BA and BPGA would be their different hydrophobic/hydrophilic moieties. Our results indicated that vaping interactions vary depending on the surface pressure, which suggests a role of the lateral packing properties. To investigate this at molecular level, we performed all-atom molecular dynamics simulations using our well-validated simulation approach that captures the physics of the air–water interface.^32,33,55,56^ We simulated the SLS composition of monolayers with two concentrations of BA and BPGA and across a range of compression states. Fig. 2A shows snapshots of the SLS monolayers containing a higher concentration of BA and BPGA at three selected area per lipid (APL, Å^2^) values. APL reflects different surface pressures, *i.e.* the lower the APL, the higher the surface pressure. These snapshots determine the preferential partition properties of the vaping chemicals in a SLS monolayer. Overall, BPGA shows a high partitioning preference to the nonpolar lipid acyl chain region at all APL values, whereas some BA always remains in the aqueous phase. Curiously, at the higher concentration, BPGA is not very soluble in the lipid phase at small APL (55 Å^2^) and forms aggregates at the lipid–air interface. Still, in a 10-fold lower concentration, BPGA is readily soluble also in the compressed monolayers, while some BA remains in the aqueous phase (not shown).

**Figure 2:**
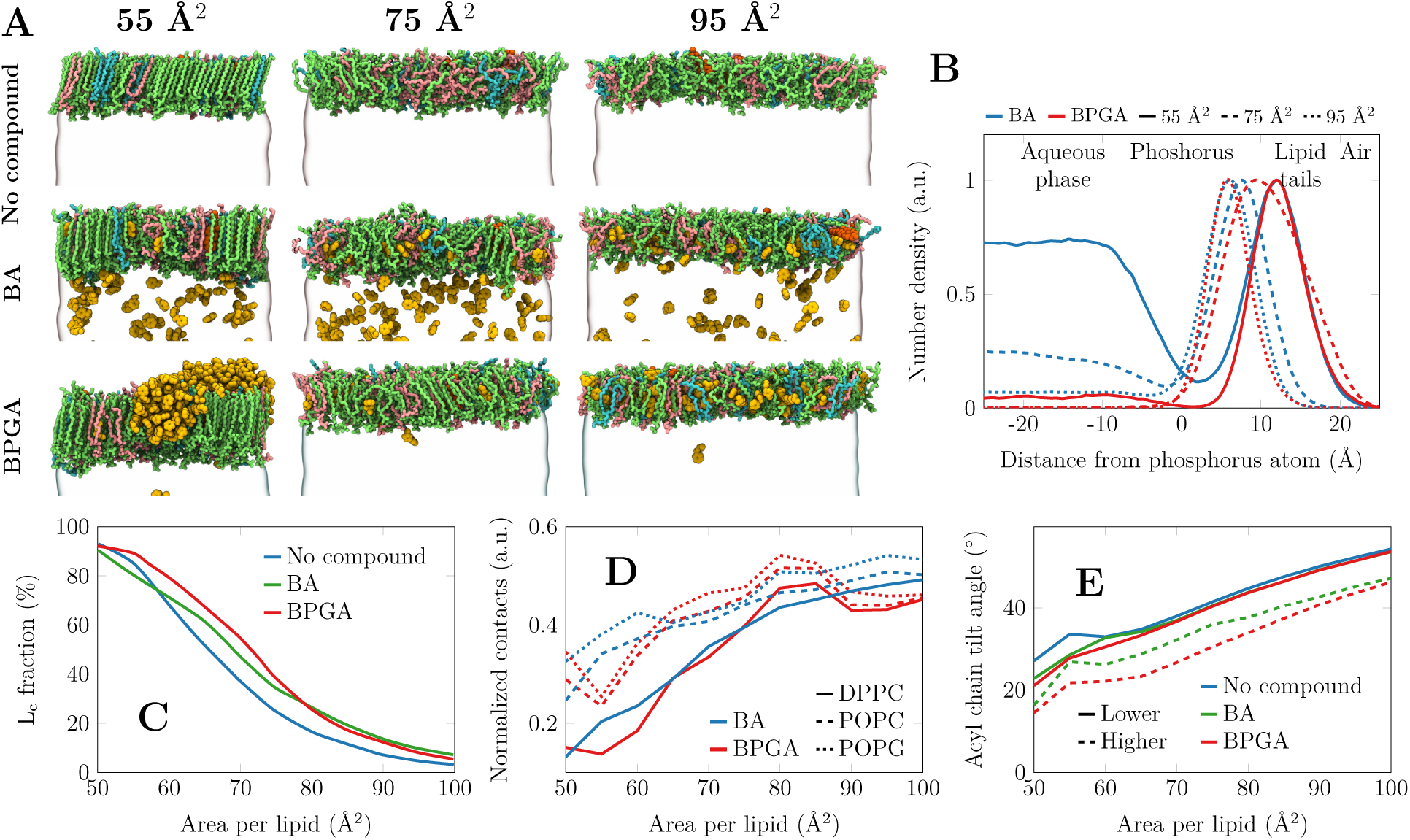
Partition preferences of the vaping chemicals into the SLS monolayer from atomistic molecular dynamics simulations. A) SLS monolayers composed of 149 lipids per monolayer, containing no vaping chemicals (top row), 320 benzaldehyde (BA) molecules (middle row), or 200 benzaldehyde propylene glycol acetal (BPGA) molecules (bottom row). The partitioning tendency at 10-fold smaller compound concentrations was similar (not shown), yet there were not enough vaping chemical molecules for aggregation. The final structures of simulations at three areas per lipid are shown. DPPC, POPC, POPG and cholesterol are depicted in green, pink, cyan, and orange respectively, whereas the vaping compounds are shown in yellow. Water is shown as a transparent surface and all hydrogens are omitted for clarity. B) The density profiles of BA and BPGA across the lipid monolayer at the air–water interface. Data are shown at three compression states also visualized on the left of the figure. The density profiles are normalized so that their maxima is set to 1. C) The fraction of acyl chains and cholesterol molecules that are tightly-packed, hence resembling the L_c_ phase. D) The interaction preference of BA and BPGA with different lipids. As the lipid moieties are present at different amounts, the contacts are normalized by the number of possible interactions. E) The tilt angle of the phospholipid acyl chains with no vaping chemicals as well as with lower and higher concentrations of either BA or BPGA.

The partitioning of vaping chemicals is quantified by density profiles in Fig. 2B. Here, the normalized (maximum=1) number density across the monolayer normal is shown. The curves confirm the visual observation from Fig. 2; BPGA always partitions to the lipid phase, whereas a substantial fraction of BA remains in the aqueous phase. In the compressed monolayer with an APL of 55 Å^2^, BPGA resides in the acyl chain region and is depleted from the polar head group region, whereas BA has significant populations in the acyl chain region, in water, and also in the head group region. At an intermediate APL of 75 Å^2^, BA prefers the polar interface and water, whereas BPGA resides deeper in the monolayer. In the very loosely-packed monolayer at 95 Å^2^, both chemicals reside in the head group region of the thin monolayer.

Next, we looked into the selective interactions of BA and BPGA with different lipid species by analyzing the vaping chemical–lipid contacts from the simulations. As demonstrated in Fig. 2D, both vaping compounds show no specificity toward any lipid type at large APLs, *i.e.* in the loosely-packed monolayer. However, upon compression both BA and BPGA demonstrate more interactions with the lipids that have unsaturated acyl chains, namely POPC and POPG in the SLS mixture. This result suggests that the compounds are excluded from the tightly-packed DPPC acyl chains and do not significantly perturb their adaptation into the L_c_-like phase. This presence of BA and more so of BPGA in the monolayer and especially among the unsaturated acyl chains naturally has implications on its structure.

Firstly, the presence of BA and BPGA lead to tighter monolayer packing. The chemicals residing among the unsaturated acyl chains at a constant area lead to a larger amount of lipid chains being assigned with the L_c_-like packing. This effect is demonstrated in Fig. 2C. At large APLs, essentially no chains are packed and the chemicals have little effect. At small APLs, on the other hand, essentially all acyl chains are packed tightly despite the presence or absence of BA or BPGA. At intermediate areas, on the other hand, both BA and BPGA promote lipid packing, and at in the range from 55 Å^2^ to 75 Å^2^ the effect of BPGA is more significant.

One notable effect of BA and BPGA at high compression is that they decrease the average lipid tilt. As demonstrated in Fig. 2E, in the compressed monolayer and in the absence of vaping chemicals, lipid acyl chains adapt a conformation with a typical L_c_-like tilt of *≈*25*^◦^* observed in experiments^57^ as well as in previous simulations of compressed lipid monolayers.^32,56^ However, even when a small amount of BA or BPGA are added, the tilt angle in the compressed monolayer decreases by *≈* 5*^◦^*. With a larger concentration of vaping chemicals interacting with the monolayer, the acyl chain tilt angles are decreased across all compression states by *≈* 10*^◦^*.

### BA and BPGA Interactions with Surfactant Lipids under Physiological Dynamics do not Significantly Influence Biophysical Function

To confirm physiological relevancy of interactions seen with SLS on the LBT, the effect of BA and BPGA were tested in physiological compression–expansion cycles on the CSD surfactometer. As seen in Figs. 3A–C, minimal differences were observed between the shapes of the *γ − A* iso-cycles of each condition. This could be confirmed after observing the lack of significance between groups for all three functional parameters (*γ*_min_, global compressibility and hysteresis) during both first and last cycles (Figs. 3D–G). Overall, the minimal loss of BA and BPGA surface-active properties under physiological dynamics indicates little interaction between SLS and BA or BPGA. This was a rather surprising result, and we thus proceeded to validate it by assessing interfacial behaviour of BA and BPGA in physiological compression– expansions in a dose-dependent manner. Each chemical surface tension was measured on the CSD in increasing concentrations **(Supplementary Fig. S3)**. 1 mg/ml BPGA had higher surface activity than BA at the same concentration, reducing surface tension from a mean of 72 mN/m to 48 mN/m, rather than 67 mN/m seen with BA. Increasing concentrations of both chemicals consistently decreased surface tension, apart from 1 g/mL BPGA, which sank immediately due to high density. These results imply that BA and BPGA remain surface-active at the air–liquid interface over multiple cycles and—considering the above results after statistical analysis (Figs. 3D,F, and G)—whilst at the interface interfere minimally with its surface tension.

**Figure 3:**
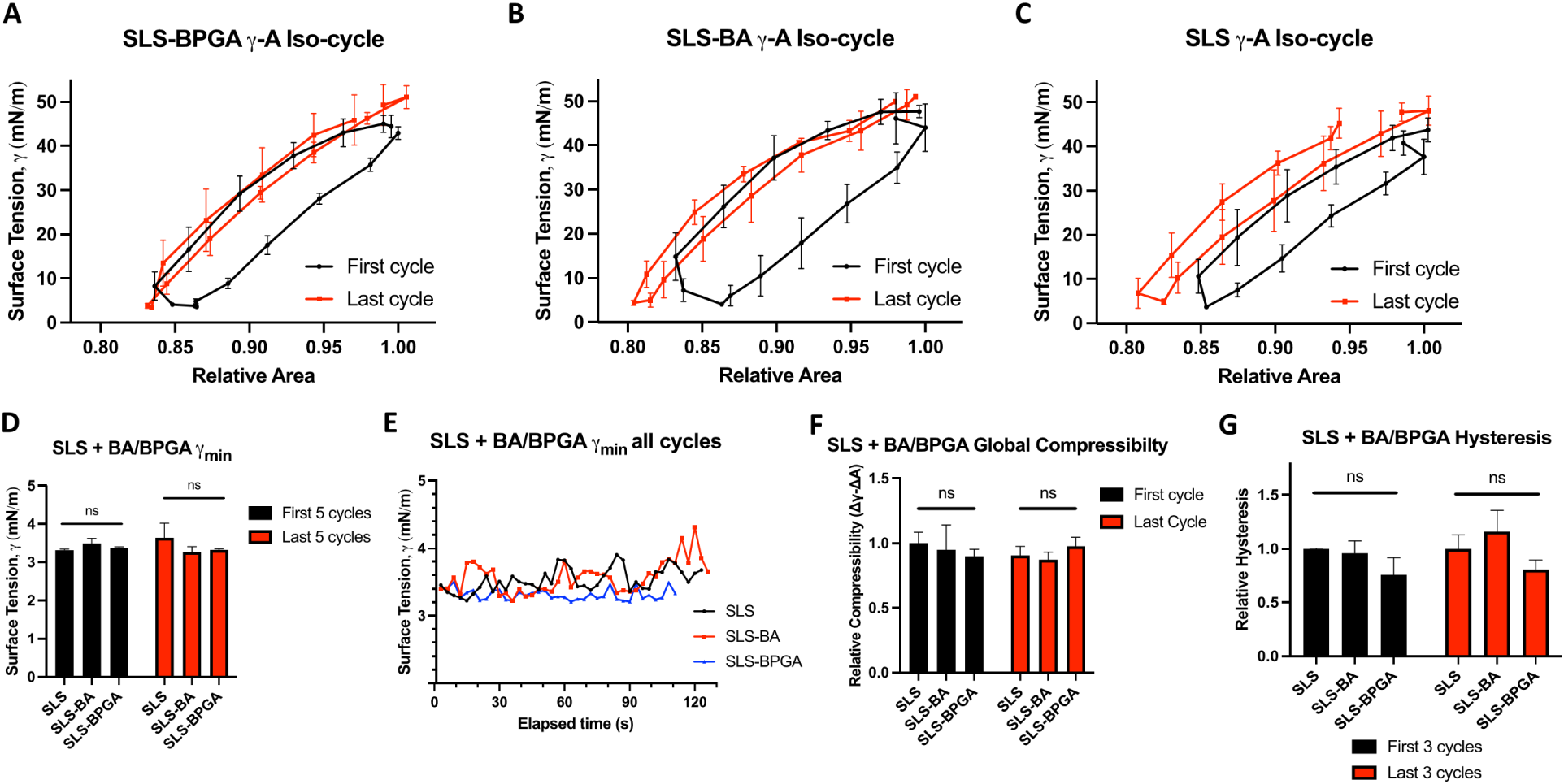
Physiological compression–expansion cycles for SLS and vaping components. Results from CSD cycles run at 20 cycles per minute for two minutes. A) SLS alone, first and last iso-cycles. B) SLS with BA, first and last iso-cycles. C) SLS with BPGA, first and last iso-cycles. BA and BPGA were added at a 1:10 molar ratio to surfactant. D) Comparison of the mean *γ*_min_ of the first and last 5 cycles for each condition. E) Temporal evolution of *γ*_min_ covering all cycles of each condition represented. F) Relative global compressibility of the first and last cycles, relative to SLS first cycle. G) Mean hysteresis of the first and last three cycles, relative to SLS first cycles. N=3 individual replicates for each condition. Significance values represent results from a Two-Way ANOVA; “ns”: not significant (p*>*0.05).

### BA and BPGA alter the Biophysical Function of Alveofact at Physiological Compression-Expansion Rates

Our previous results indicate that physiological dynamics induce loss of lipid interactions with BA and BPGA. Still to be investigated is whether the addition of surface-active surfactant proteins would influence the chemical interactions of vaping components at the interface, and whether this would induce disruption in the biophysical function of the surfactant.

Alveofact is a commercial bovine-derived clinical surfactant containing the hydrophobic surfactant proteins SP-B and SP-C.^45^ In attempts to assess whether the addition of SP-B and SP-C would result in additional molecular interactions with BA and BPGA from those observed with surfactant lipids, Alveofact was tested on the LBT. In contrast to SLS **(Supplementary Fig. S4A)**, Alveofact would not reach the standard Π_max_ of *>* 70 mN/m,^36^ rather plateauing at 50 mN/m. This limit was confirmed by increasing the molar concentration of Alveofact up to 5-fold **(Supplementary Fig. S4B)**. Therefore, LBT measurements would not be informative of changes to biophysical properties at physiologically relevant high surface pressures.

For this reason, Alveofact experiments progressed to the CSD, where it was theorised that fast physiological compression–expansion rates of 20 cycles per minute^50^ would force rapid lateral self-assembly, reducing the accumulation of surfactant constituents into the 3D reservoir and thus enabling the achievement of *γ*_min_. This would allow the observation of physiologically relevant molecular interactions. To this end, the effect of vaping chemicals on the biophysical function of Alveofact were evaluated over 40 physiological compression– expansion cycles **(Supplementary Fig. S5)**. BA and BPGA caused visible alterations to Alveofact *γ − A* iso-cycles extracted from cycle data (Figs. 4A–C). While Alveofact iso-cycles remain stable throughout multiple cycles, BA and BPGA addition seem to induce iso-cycle deformation, especially during the first iso-cycle. Interestingly, these observations exhibit interference of surface tension and area change and thereby biophysical properties.

**Figure 4:**
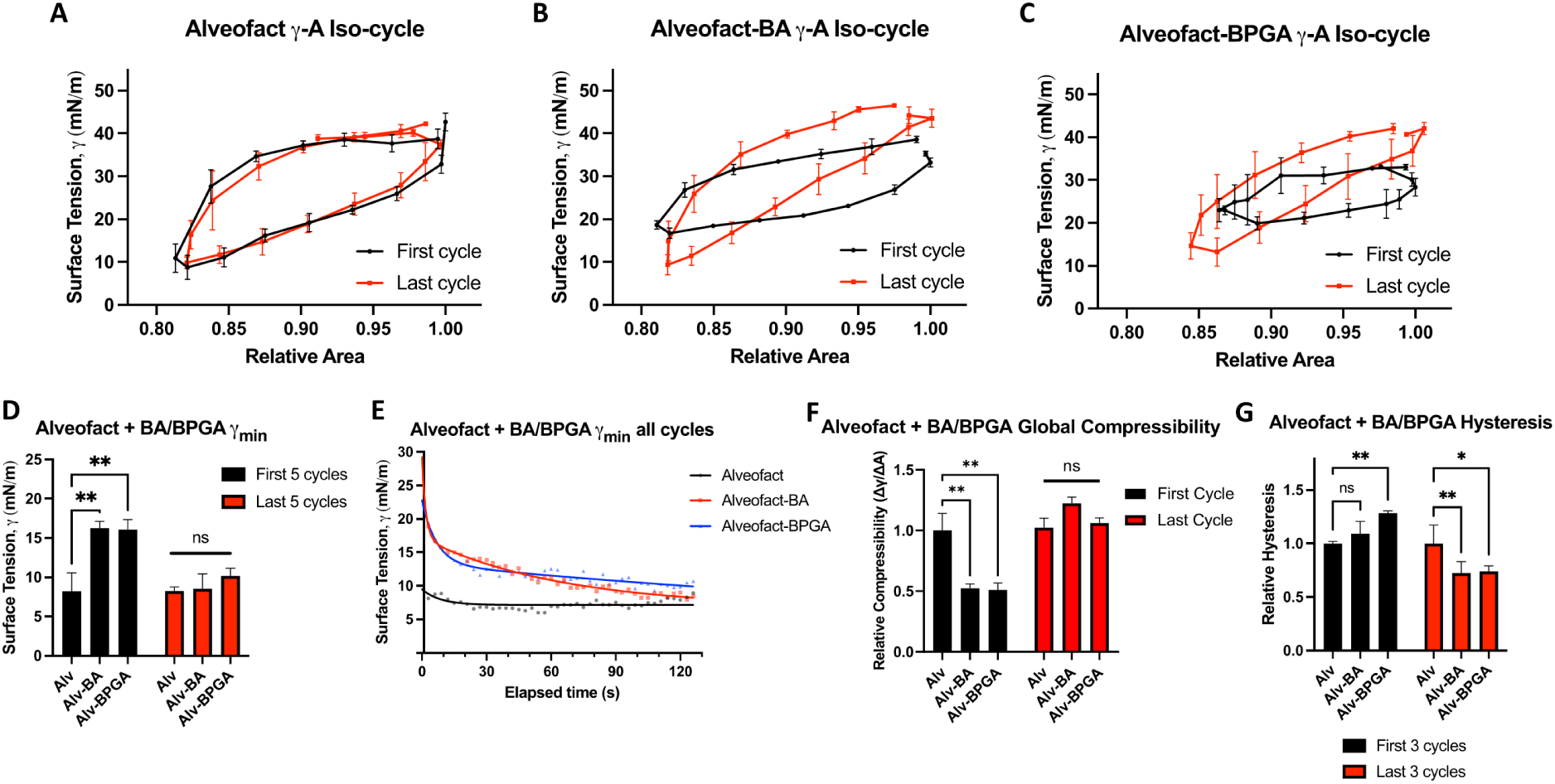
Physiological compression–expansion cycles for Alveofact and vaping chemicals. Results from CSD cycles run at 20 cycles per minute for two minutes. A) Alveofact alone, first and last *γ − A* iso-cycles. B) Alveofact with BA, first and last *γ – A* iso-cycles. C) Alveofact with BPGA, first and last *γ − A* iso-cycles. BA and BPGA were added in a 1:10 molar ratio to the surfactant. D) Comparison of mean *γ*_min_ of the first and last 5 cycles for each condition. E) Temporal evolution of *γ*_min_ over all cycles for each condition. Alveofact alone is fitted to a mono-exponential decay curve. Alveofact–BA and Alveofact–BPGA are fitted to bi-exponential decay curves. F) Global compressibility of the first and last cycles relative to the first cycle of Alveofact. G) Mean hysteresis of the first three and last three cycles, relative to the first three cycles for Alveofact. N=3 individual replicates for each condition. Significance values represent results from a Two-Way ANOVA; “ns”: not significant (p*>*0.05), *p*<*0.05, **p*<*0.01.

To quantify iso-cycle alterations, firstly, *γ*_min_ were extracted (Figs. 4D–E). In the first cycles, there were significant increases in the mean surface tension from 8 mN/m with Alveofact alone to 16 mN/m with either BA or BPGA included (Fig. 4D). This effect was lost during the last five cycles, during which BA- and BPGA-containing mixtures have similar *γ*_min_ values as Alveofact alone. This trend was reflected in exponential decay curves representing kinetics of all cycle *γ*_min_ values (Fig. 4E). A shorter decay half-life indicates faster reduction in *γ*_min_, *i.e.*, an improvement in biophysical function. While Alveofact alone fits a mono-exponential curve and quickly plateaus to a constant *γ*_min_ of *≈*7 mN/m, indicating rapid monophasic improvement, the addition of BA or BPGA causes a shift to bi-exponential curves which plateau at higher surface tensions. This infers that the rapid decrease is coupled with a long-term interference of biophysical function, with BPGA inducing the most persisting disruption to *γ*_min_ with a second phase half-life five times slower than that of BA (253.2 s compared to 44.52 s). In brief, the experiments suggest that BA and BPGA interfere with the biophysical function of the surfactant by interacting with the surfactant proteins SP-B and SP-C in physiological compression–expansions over time scales of minutes.

To assess the effects of protein–vaping chemical interactions on film compaction and elasticity, global compressibility modulus was calculated *via* the iso-cycle slope.^49^ BA and BPGA significantly reduce the global compressibility modulus *κ* of Alveofact during the first cycle by *≈*50%, although this is restored during the last cycles (Fig. 4F). Therefore, protein–vaping chemical interactions initially decrease lateral film compaction, although this effect is not sustained under physiological dynamics. Finally, to determine if BA and BPGA induce alterations in membrane organisation over multiple physiological compression–expansion cycles, Alveofact hysteresis was extracted from the first and last *γ − A* iso-cycles. Hysteresis of the first five cycles increased with BA and BPGA addition, although BPGA imposed the only significant change (Fig. 4G). Interestingly, during the last 5 cycles, BA and BPGA both significantly *decrease* Alveofact hysteresis. Protein–vaping chemical interactions thereby induce persistent changes to film organisation. BA and BPGA induce material loss in the first cycles, which in turn presents a more refined—yet possibly less functional—monolayer by the last cycles compared to Alveofact alone.

### BA and BPGA Interact with Surfactant Properties in the SLS Monolayer

Our experiments on Alveofact suggested that BA and BPGA could interact with the surfactant proteins SP-B and SP-C, leading to persistent negative impacts on the parameters characterizing the biophysical function of the surfactant—namely *γ*_min_ and hysteresis under physiological compression–expansion dynamics. To validate this hypothesis, we performed additional atomistic molecular dynamics simulations of surfactant monolayers containing either SP-B or SP-C. Moreover, these simulations followed the experiments, as BA or BPGA were first allowed to interact with the monolayer, after which it was compressed from a large area per lipid of 110 Å^2^ (Π *≈* 0 *mN/m*) to a small one of 55 Å^2^ (Π *≈* 70 *mN/m*) during the course of a 2 µs-long simulation. Examples of the simulation systems with BPGA are shown on the left panel of Fig. 5.

**Figure 5:**
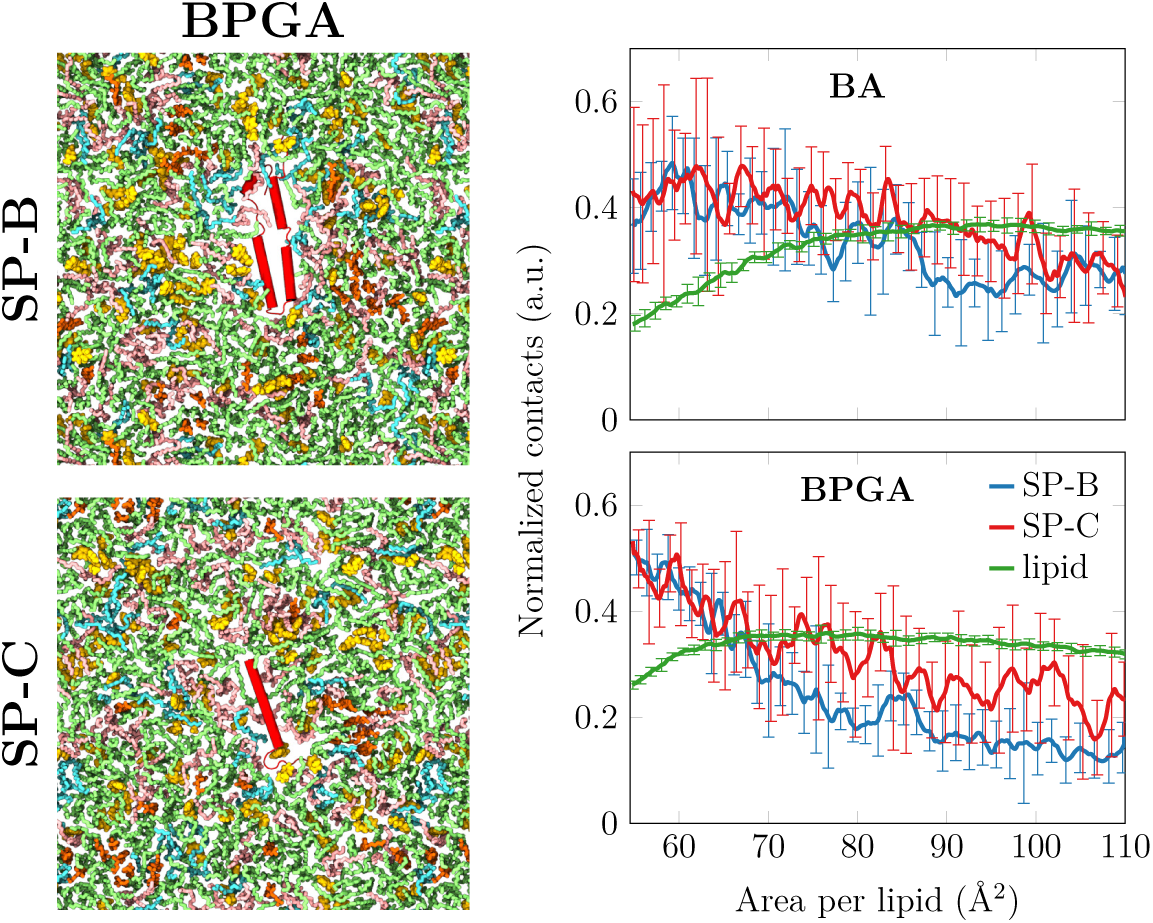
Interaction of vaping chemicals with hydrophobic surfactant proteins SP-B and SP-C. Left: Snapshots of the simulation system containing one copy of either SP-B (top) or SP-C (bottom). The shown simulations also contain BPGA, shown in yellow. Proteins are shown in red, DPPC in green, POPC in pink, POPG in blue, and cholesterol in orange. Hydrogens and waters are omitted for clarity. Right: Interactions of BA and BPGA with the surfactant lipids and proteins as a function of monolayer area. The contact numbers are normalized by the number of possible interactions in the two groups included in the analyses. For lipids, only the curve from the simulation containing SP-B is shown since the result with SP-C is essentially identical. The error bars show the difference between the data calculated from the two replica simulations.

We calculated the number of contacts between the vaping chemicals and the lipids or the protein, as these values characterize their preferential interaction partners at different compression states. As demonstrated on the right panel of Fig. 5, the interactions with lipids are somewhat sensitive to the compression state. At low areas per lipid, more BA remains in the aqueous phase, leading to a steady decrease of contacts below *≈* 80 Å^2^. The same exclusion takes place with BPGA, but to a smaller extent due to its larger partitioning preference towards the hydrophobic acyl chain region (Fig. 2A,B).

Surprisingly, the interaction of BA and BPGA with the surfactant proteins demonstrates a different trend. When the monolayer is compressed, both BA and BPGA accumulate near the proteins. In this state, interactions with SP-B and SP-C are equally likely. However, these contact numbers are normalized by the possible interaction partners, signaling that the larger SP-B interacts overall with more vaping chemicals. Overall, BPGA demonstrates more interactions with the proteins in the compressed monolayer. Therefore, our atomistic simulations confirm that both BA and BPGA indeed interact with the surfactant proteins especially in the physiologically relevant compressed surfactant, leading to the compromised biophysical function of Alveofact observed in the experiments.

## Discussion

In this study, we evaluated the effects of the highly hydrophobic e-liquid by-product, BPGA, and its hydrophobic precursor, the flavouring aldehyde BA, on the biophysical function of surfactan tunder dynamic conditions. Our aim was to assess the role of common vaping flavourings in surfactant dysfunction. It was hypothesised that these components may affect the surfactant due to their hydrophobic nature, thereby enabling interactions with surfactant molecules at the air–liquid interface. BPGA, as the more hydrophobic molecule, was expected to have a stronger impact on surfactant biophysical function than BA.^44^ Fine interactions between surfactant lipids with both BA and BPGA were confirmed at quasi-static compression–expansion rates employing a lipid-only SLS model (Fig. 1). Our atomistic molecular dynamics simulations also demonstrated the partitioning of both BA and BPGA into the SLS monolayer at all compression states, with BPGA exclusively residing in the acyl chain region at high surface pressures.

BA and BPGA induced significant reduction of initial maximum surface pressures, likely explained by the additional negative impact on compressibility, as both chemicals impaired lateral compaction at low surface pressures and maximum surface pressure, where lipids did not self-assemble into the gel phase. Furthermore, initial SLS hysteresis was significantly reduced. While this does not directly indicate loss of surface-active material, it may imply a lack of film refinement, preventing optimisation of monolayer arrangement.^28^ The effect of BA and BPGA observed on the compressibility modulus at a surface pressure of *≈*20 mN/m (Fig. 1D) coincides with the region where ordered L_c_-like domains start to form in the surfactant.^32^ With BA or BPGA present in the membrane, more chains are involved in these domains (Fig. 2C), thus decreasing the overall lipid packing and thus the compressibility modulus. At high pressures, the compressibility modulus is increased by the presence of BPGA (Fig. 1D). This coincides with the aggregation of BPGA molecules in the monolayer (Fig. 2A) at this pressure range, and these clusters could resist compression.

Notably, the significant effects on Π_max_ and hysteresis were either lost or considerably reduced over the 10 cycles. A potential explanation may be the “squeeze-out” hypothesis, where the surfactant lipids with saturated acyl chains, *i.e.*, DPPC, force other molecules out from the monolayer at high compressions.^58^ This would usually cause material rearrangement into the 3D structure reservoir, however, in a lipid-only model, the lack of proteins limits the stability of unsaturated phospholipid sublayer formation, which increases collapse and therefore loss of chemicals directly into the subphase. Unfortunately, the sizes of atomistic simulation system sizes are too small to observe this “squeeze-out”. Still, the decrease in surface pressure during the first cycle could well result in the excessive structural perturbation summarized in Fig. 2; the vaping chemicals interact preferably with the unsaturated lipid chains and thus affect the phase behavior of the surfactant, including the decrease in lipid tilt characteristic for the L_c_ phase required for reaching the physiological Π_max_.

The fine surfactant lipid–chemical interactions were then proved insignificant under physiological compression–expansion dynamics, as SLS CSD results presented no significant differences in any of the three functional parameters (Fig. 3). Therefore, BA and BPGA are likely not able to significantly disrupt pulmonary surfactant biophysical function *via* lipid interactions. A probable reason is immediate chemical loss from the interface imposed by rapid compression rates, forcing abrupt maximum lateral compaction, after it was observed that BA and BPGA would normally impose surface tension reduction under these conditions **(Supplementary Fig. S3**, Fast lateral compression has been observed to eventually force components out of the monolayer towards the linked bilayers beneath, ^30,32,53^ which seems to be a very plausible mechanism with vaping components too.

Once lipid interactions indicated physiological surfactant dysfunction insignificance, the investigation turned to potential interactions between BA and BPGA and the surface-active surfactant proteins SP-B and SP-C, by employing the clinical surfactant Alveofact.^45^ Unfortunately, attempts at fine interaction studies with Alveofact at quasi-static rates **(Supplementary Fig. S4)** were identified as abnormal due to the knowledge that Alveofact must reach *γ*_min_ below 2 mN/m to be an effective surfactant substitute in premature neonates.^20,45^ Additionally, similar behaviour of Alveofact has previously been reported on the LBT.^59^ The slow quasi-static compression rates likely enable SP-B and SP-C to facilitate extensive monolayer folding, or 2D to 3D transition, reducing maximum lateral lipid compaction by Π_max_.^24,34^ Physiological compression–expansion rates provided a solution by forcing rapid lateral compaction to overcome protein-facilitated film elasticity. Under physiological dynamics, BA and BPGA were proven to induce persistent disruption of the biophysical function of Alveofact *via* stable vaping chemical–protein interactions, after long-term interference of both *γ*_min_ and hysteresis were observed (Figs. 4E,G). Essential protein involvement was deduced as protein content is the most prominent difference between SLS and Alveofact, and therefore the most plausible reason for differences observed at physiological dynamics. Protein interactions are likely more stable than with lipids due to the additional potential to form intermolecular forces with charged protein groups.^33,50^ Indeed, such interactions were also observed in our atomistic molecular dynamics simulations of protein-containing monolayers under dynamic compression. Whereas compression leads on average to fewer lipid–vaping chemical interactions, the number of surfactant protein–vaping chemical interactions actually increased. This suggests that BA and especially BPGA can perturb the functions of the proteins at this physiologically relevant low-tension state. Vaping chemical interactions with SP-B and SP-C may also provide reasoning for the initial loss of effect on all biophysical function parameters (Fig. 4), which is especially evident in the rapid first phase of *γ*_min_ decrease (Fig. 4E). This first phase with higher biophysical impact may represent pre-optimisation of film arrangement, where BA and BPGA have not yet fully associated with the surfactant proteins, at which point they would be able to transiently enter the 3D reservoir along with SP-B and SP-C throughout multiple maximum compressions.^24,34^ This would avoid permanent “squeeze-out” from the surfactant film, while also preventing chemicals interfering with surface-active surfactant lipid self-assembly at points of high lateral compaction. That said, it is likely that much of the vaping chemical is still being lost to the subphase from the 3D reservoir, as there remains a downward tendency in *γ*_min_ after the initial first phase of decline (Fig. **??**E). In a cellular model, this may have negative implications, as once past the surfactant film, chemicals can freely diffuse through the alveolar liquid to reach the epithelium, where they may induce cytotoxicity. ^60^ An inflamed alveolar epithelium can in turn lead to further surfactant disruption *via* exposure to foreign molecules.^61^

Throughout the study, it was unexpectedly observed that BA had a similar effect to BPGA on surfactant biophysical function, especially after results demonstrated BPGA has greater surface activity **(Supplementary Fig. S3)**. However, this did not translate into the context of surfactant interactions, with results presenting similar significance. Although BPGA did cause a longer-term disruption of Alveofact *γ*_min_ (Fig. 4E), the proposal that BA chemical alone has potential to disrupt surfactant function, thereby possible EVALI development, is concerning. This is because BA is a highly popular vaping flavouring, present in approximately 75%^41^ of flavoured e-liquids popular in the younger population. Furthermore, BA is far from the only flavouring aldehyde used in e-liquids that has acetal by-products. Other commonly used ones include vanillin and cinnamaldehyde.^62^ In future, a larger range of aldehyde flavourings should be tested to widen result relevancy.

## Conclusions

This study highlights the need for further investigation into the safety of aldehyde flavourings and other food-grade flavourings that are assumed safe for inhalation by the vaping industry prior to comprehensive long-term testing. This is after findings indicate the flavouring aldehyde BA and its respiratory irritant by-product, BPGA have potential to disrupt surfactant function. Results also suggest that upon continuous breathing, vaping chemicals will be dragging surfactant components down to the aqueous subphase. This suggest a high likelihood for the chemicals to reach the following layer of the alveolar capillary barrier, *i.e.*, the alveolar epithelium, and eventually, inducing cell toxicity. This research may prove important information for policy makers in regards to prevent the use and abuse of these common additive in e-cigarettes. Most important, this study flags the importance of investigating the *de novo* production of by-products and not only the main ingredients. Caution should be placed not only with the large numbers of adolescents exposed to and addicted to vaping products, but also adults who aim to benefit from a safer alternative to cigarettes. In the meantime, e-cigarette use, especially those containing flavourings, should be discouraged in the younger population and regulations revised while product safety is being thoroughly examined.

## Experimental

### Surfactant Models

The SLS mixture was designed and developed from lipidomic analysis and literature review^27,46^ and the final lipid used and ratios aim at keeping the saturation, unsaturation, charged lipids and neutral lipid composition of lung surfactant: 68:20:10:2 weighted ratio of DPPC:POPC:POPG:cholesterol. ^32,33,63^ All lipids sourced from Avanti Polar Lipids (USA), and dissolved to 1 mg/ml in 2:1 chloroform–methanol (Sigma-Aldrich, Germany). The bovine-derived clinical surfactant Alveofact®(45 mg/ml, Lyomark Pharma, Germany) was diluted in 0.9% saline (pH 5.8, Sigma-Aldrich) to 1 mg/ml. The experimental chemical to lipid ratio used was high above that of one e-liquid dose.^41–43^ However, this may represent levels resulting from long-term vaping, assuming chemical accumulation at the alveolar interface, as previously suggested for VEA.^14^

### Compression–Expansion Models

#### Langmuir–Blodgett Trough (LBT)

The LBT (NIMA Technology Ltd., England) was custom-designed with a continuously enclosed Teflon-vitrified coated ribbon replacing classical barriers and filled with 0.9% saline (pH 5.8) at 25*^◦^*C. Surface pressure was quantified *via* a Wilhelmy cellulose plate to produce surface pressure–area (Π *− A*) isotherms **(Supplementary Fig. S6**. The LBT was operated *via* NIMA software. To ensure LBT accuracy, DPPC Π *− A* isotherms were reproduced regularly as control according to literature.^30^ As an additional control, before each test a saline (Π *− A*) isotherm was produced to verify constant surface pressure of 0 mN/m, *i.e.* the surface pressure of water. Thereafter, 20 µL SLS was deposited at the air–liquid interface with a Hamilton gas-tight syringe (Hamilton Company, U.S.A), before being left for five minutes to equilibrate and self-assemble prior to initiating ten compression–expansion cycles at 150 cm^2^/min, moving between 215 cm^2^ area to 56 cm^2^. Alternatively, after five minutes, BA (no.418099, Sigma-Aldrich) or BPGA (no.W213000, Sigma-Aldrich), diluted to 1 mg/ml in 2:1 chloroform–methanol, were added at a chemical to lipid molar ratio of 1:10 prior to iso-cycling. Up to three technical repeats were collected for each condition. Data were extracted from the NIMA software, which records the surface area (A, cm^2^) and surface pressure (Π, mN/m). Changes in area were recorded as relative changes form the total surface area that our trough has *≈*230 cm^2^ to enable isotherm comparisons from other troughs.

#### Constrained Sessile Drop (CSD) Surfactometer

A custom-designed constrained sessile drop system was employed. This system uses elements of the CSD surfactometer from Krüss and its software (Drop Shape Analyser). The Krüss Advance Software, which calculates surface tension *via* the contact angle between the sessile drop and pedestal (Supplementary Fig. S6B), allowing the production of *γ − A* iso-cycles.

The custom-built pedestal made of stainless steel was adapted to fit the equipment (Krüss, Germany). A microsyringe (ILS, Germany) was used connected to a stepper motor computer-controlled system to produce the fine regulated physiological compression–expansion cycles. Drop formation, recording and analysis were performed *via* recordings from an UI-3060CP Rev. 2 camera (IDS, Germany). Drops consisted of 0.9% saline with a volume of 12–14 µL. Oscillations were designed to mimic physiological conditions, meaning a 20% reduction of surface area and a rate of up to 20 cycles per minute. ^50,64^ Each replicate was run for 120 s, with a recording rate of 5 frames per second. Cycles were completed at room temperature (25*^◦^*C). For each replicate, saline was run alone to ensure approximate 72 mN/m surface tension before adding 1 µL SLS or 3 µL Alveofact to the air–liquid interface with a Hamilton pipette. These quantities were determined by selecting the volume necessary to reach the *γ*_min_ possible for each model (approximately 3 mN/m and 7 mN/m, respectively). When required, 1 mg/ml BA or BPGA were added immediately after surfactant at a molar ratio of 1:10. Drops were left for five minutes prior to initiating compression–expansion cycles. Three individual technical replicates were gathered for each condition.

### Parameters Characterising the Biophysical Function of the Surfactant

#### Minimum Surface Tension (*γ*_min_)

Surface pressure is a direct correspondent of surface tension (both have units of mN/m), *via* the equation Surface pressure (Π) = Surface tension of water (*γ*_0_) - Surface tension (*γ*). The surface tension of water at a 20*^◦^*C air–water interface is 72.8 mN/m, ^65^ meaning that at the minimum surface tension (*γ*_min_) of 0 mN/m, surface pressure is at its maximum (Π_max_) of *≈*72 mN/m. Changes in (*γ*_min_) and or (Π_max_) inform of surfactant functionality and stability.^66^ (*γ*_min_) or (Π_max_) for each cycle were extracted from exported CSD or LBT data, respectively.

#### Compressibility modulus (*κ*)

In the surfactant context, compressibility modulus (*κ*, mN/m) is a measure of resistance to lateral compression of a lipid film. Higher compressibility modulus corresponds to a compact lateral self-assembly, thus low film elasticity.^67^ It is defined as

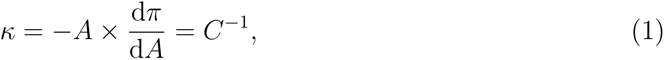

with *C* being the compressibility sometimes reported in monolayer studies.

This theory was applied to the first LBT Π *− A* isotherms using Origin software (Origin-Lab, USA). Savitsky–Golay 9-point smoothing was then employed for improved visualisation. For CSD data, the number of data points in each iso-cycle were insufficient for comprehensive compressibility moduli, rather a global compressibility was estimated by calculating the slope between minimum to maximum iso-cycle points (Δ*γ/*Δ*A*).^68^

#### Hysteresis

High hysteresis values describe a refinement or loss of material between compression and expansion, altering lateral lipid organisation.^28^ In this study, hysteresis was represented by the difference in area (Δ*A*, cm^2^) between compression and expansion isotherms at half surface pressure (LBT) or tension (CSD). For LBT data, hysteresis from the first and last cycles were collected. Due to the large cycle numbers in CSD data, this dataset increased to the mean of the first and last three cycles.

### Atomistic Molecular Dynamics Simulations

We performed two sets of atomistic molecular dynamics simulations to characterize the interaction of the vaping chemicals on pulmonary surfactant monolayers.

#### Static Simulations of Protein-free Monolayers

First, protein-free surfactant monolayers were simulated at 11 fixed areas per lipid, ranging from 50 Å^2^ to 100 Å^2^ with 5 Å^2^ intervals and thus covering the physiologically relevant compression states. The simulation contained two monolayers separated on one side by a slab of *≈*27,040 water molecules (*≈*91 per lipid) with *≈*150 mM NaCl and on the other side by a large slab of vacuum. The two monolayers each contained a total of 149 lipids with molar ratios of 68/20/10/2 of DPPC/POPC/POPG/cholesterol, *i.e.* in accord with the experimental SLS mixture. The simulations were performed with either low or high concentrations of BA (32 or 320 molecules) or BPGA (20 or 200 molecles). A vaping chemical-free system was simulated as a control. All simulations were 1 µs long.

The amount of L_c_-like packing was characterized by clustering the 10^th^ carbon atoms along the acyl chains of phospholipids and the C14 atom of cholesterol in the plane using the DBSCAN algorithm with a cut-off of 0.71 nm and a requirement for 6 neighbours within this cutoff.^32^ Any acyl chains or cholesterol molecule that were found to be part of a tightly-packed cluster was assigned to the L_c_ phase, and the total fraction of this phase was then averaged over time for each static simulation.

The contact preferences with lipids were calculated using `gmx mindist` tool bundled with GROMACS. Only non-hydrogen atoms were included to speed up the analysis, and the cutoff was set to 0.6 nm. All contacts were normalized based on the number of possible interaction partners.

The density profiles were calculated using `gmx density` bundled with GROMACS. The profiles were centered at the phosphate position, and all profiles were normalized to have a maximum value of 1.

The tilt of acyl chains was averaged over all phospholipids, over their two chains, and over time. The tilt angle was defined as the angle between the vector connecting the 1^st^ and 16^th^ carbons of the acyl chains and the normal to the monolayer (*z* axis).

#### Dynamic Simulations with Surfactant Proteins

Secondly, we performed dynamic simulations of the pulmonary surfactant monolayers in which the monolayer was compressed so that the APL decreased from 110 Å^2^ to 54.5 Å^2^ in the course of a 2 µs-long simulation. The monolayer composition was 60/20/10/10 of DPPC/POPC/POPG/cholesterol, following our earlier work^32,33^ and hence slightly different from the SLS mixture used in experiments and in the static simulations. Two monolayers present in the simulation system were again separated by a slab of water (38400 molecules, *≈*75 per lipid with *≈*150 mM of NaCl) on one side and vacuum on the other side (across the periodic boundary conditions). The monolayers contained either only lipids or a single copy of SP-B or SP-C each.^33^ Each system was simulated in the absence of vaping chemicals as well as in the presence of 320 molecules of BA or 200 molecules of BPGA. The simulations were performed in duplicate.

From the dynamic simulations, we analyzed the numbers of contacts between the vaping chemical and the surfactant proteins as well as the lipids. This analysis was performed over the dynamic trajectory. The two replica simulations were analyzed, and their mean values and differences were used as the reported result and its error estimate, respectively. Hydrogens were omitted from the analysis, and the numbers were normalized based on possible interaction partners present in the simulation. A cutoff of 0.6 nm was used to define a contact.

#### Force Fields and Simulation Parameters

We used the CHARMM36 model to describe the lipids^69,70^ and the 4-point OPC water to model water.^71^ This force field combination has successfully captured the interfacial physics of the water–air interface and monolayers placed thereon. ^32,55,56^ The vaping chemicals were described with the Merck molecular force field^72^ obtained from SwissParam.^73^ For proteins, the CHARMM36m force field^74^ was used with the protein models adapted from our previous work.^33^

For the static simulations with a fixed monolayer area, the simulation protocol followed our earlier work^32^ except that the vaping chemicals were originally placed in the aqueous phase. For the dynamic simulations with slowly changing monolayer area, we also followed our earlier work^33^ apart from the presence of the vaping chemicals.

Simulation inputs and outputs are openly available in the Zenodo repository at DOI: (Will be uploaded upon acceptance).

### Statistical Analysis

Where appropriate, Two-Way analysis of variance (ANOVA) tests was performed, followed by Dunnett’s multiple comparisons tests to compare every group mean to the control. Significance was determined with alpha set to 0.05. Standard error measurement was calculated and presented as error bars where suitable. All graphs and statistical analyses were produced utilising GraphPad Prism 9.1.0 Software (GraphPad, USA).

## Acknowledgement

J.B.S acknowledges support from Bill & Melinda Gates Foundation and BBSRC (Grants INV-016631 and BB/V019791/1, respectively); also from the Cancer Research UK Convergence Science PhD Scholarship to Alexia Martin and a pump-prime award from the Integrated Biological Imaging Network (IBIN), a Technology Touching Life MRC Network (MR/W024985/1). We would like to thank Lyomark Pharma GmbH and in particular Dr Daniel Kune for providing Alveofact samples. M.J. acknowledges the Emil Aaltonen foundation and the Academy of Finland (grant no. 338160) for funding. We thank CSC–IT Center for Science (Espoo, Finland) for computational resources. I.V. thanks the support granted by the Academy of Finland (projects 331349, 336234, 346135), the Sigrid Juselius Foundation, Helsinki Institute of Life Science (HiLIFE) Fellow Program, the Lundbeck Foundation, and the Human Frontier Science Program (RGP0059/2019).

## Supplementary Information

### Supplementary Methods

#### SLS LBT Temperature Experiments

SLS (outlined in main methods) was trialed on the LBT at 25*^◦^*C as well as 37*^◦^*C, maintained with a TC120 water bath (Grant Instruments, UK), where both 20 µL and 35 µL volumes of 1 mg/ml SLS were used in attempts to reach Π_max_ of 72 mN/m. SLS was deposited onto the air–liquid interface with a Hamilton gas tight syringe (Hamilton Company, U.S.A), before being left to adsorb to the interface for five minutes prior to proceeding with compression at a rate of 150 cm^2^/min, moving from maximum area 215 cm^2^ to minimum area 56 cm^2^.

#### Alveofact LBT Trials

When using Alveofact (Lyomark Pharma, Germany) on the LBT, 20 µL 1 mg/ml was initially run on the LBT with similar procedure to SLS. In attempts to reach higher surface pressures, Alveofact concentration was increased to 10 mg/ml (diluted in 0.9% saline, Sigma-Aldrich) and added in volumes of 4 µL and 10 µL (2*×* and 5*×* molar quantities, respectively).

#### Vaping Chemical Surface-activity

BA and BPGA were tested alone on the CSD model. BA (no.418099, Sigma-Aldrich, Germany) and BPGA (no.W213000, Sigma-Aldrich) were serially diluted to 1 mg/ml from 1 g/ml in factors of 10 in 2:1 chloroform-methanol (Sigma-Aldrich). After first running saline alone, 1 µL of each concentration was added to the drop and run for 60 s at 20 cycles per minute consecutively in ascending order without prior surfactant addition.

### Supplementary Results

**Figure S1:**
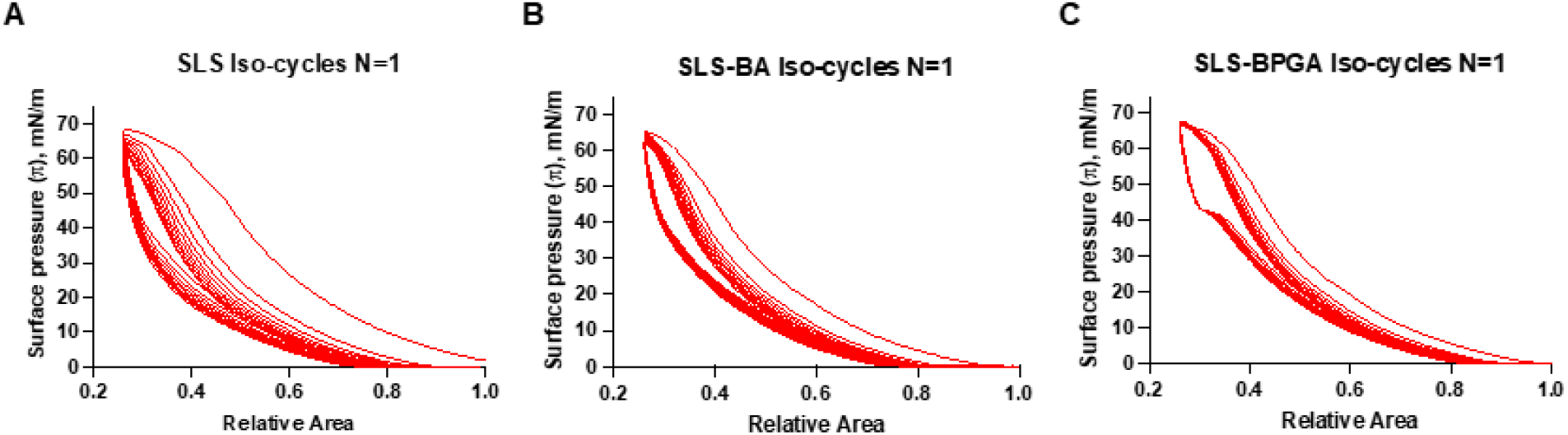
Representative Π *− A* Iso-Cycles for SLS with BA or BPGA. A) SLS alone. B) SLS with BA at a 1:10 molar ratio. C) SLS with BPGA at a 1:10 molar ratio. Cycles were completed on a saline subphase at 25*^◦^*C with a compression rate of 150 cm^2^/min over ten cycles. Although up to N=3 replicates were collected, N=1 from each condition is presented for clarity.

**Figure S2:**
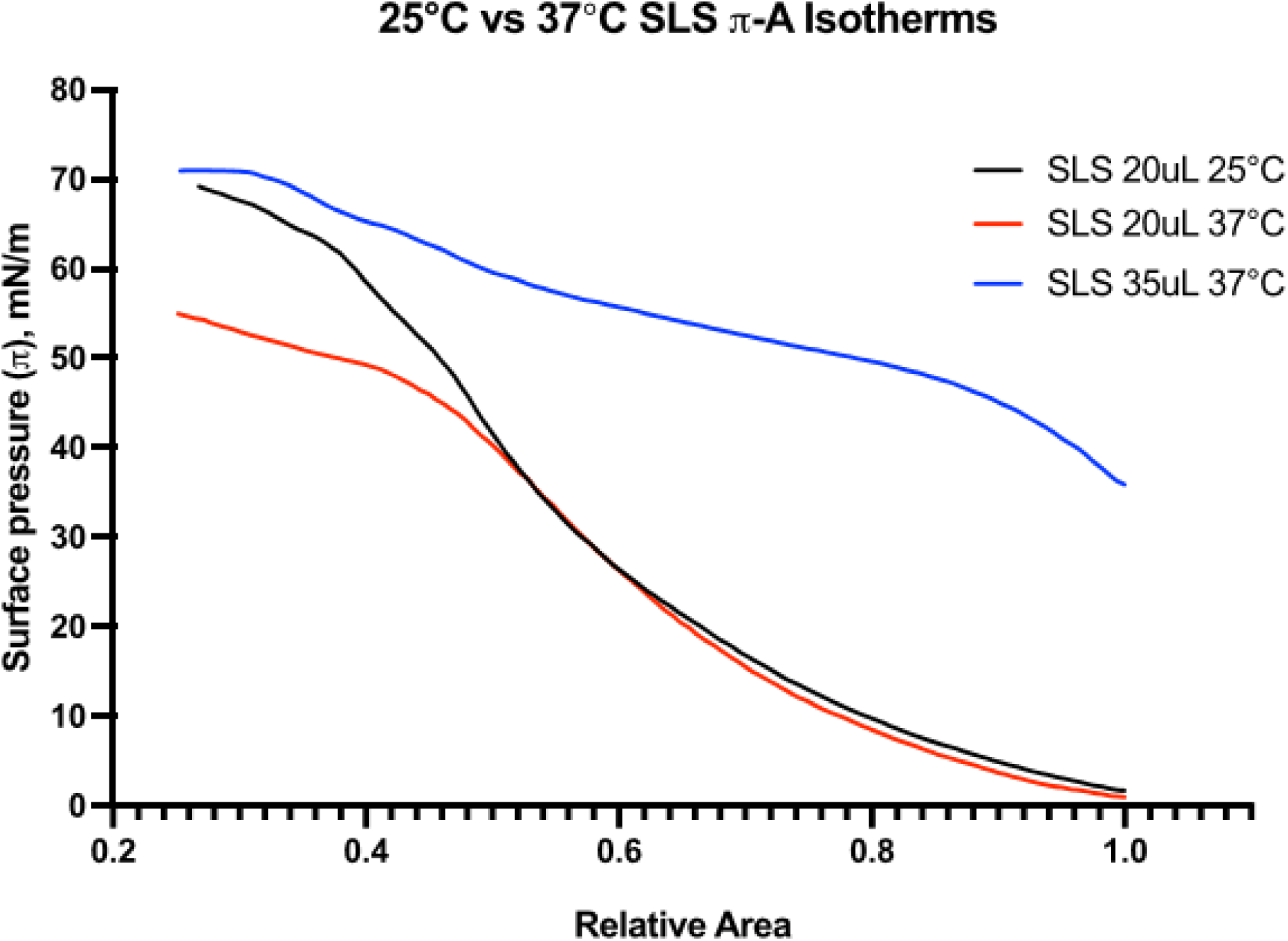
SLS LBT temperature experiments. A 25*^◦^*C Π *− A* isotherm is compared to an isotherm at 37*^◦^* also produced with 20 µL 1 mg/ml SLS, and an isotherm produced with 35 µL volume. at a 1:10 molar ratio. Isotherms were completed on a saline subphase at 25*^◦^* with a compression rate of Π *− A*. N=3 for each condition.

**Figure S3:**
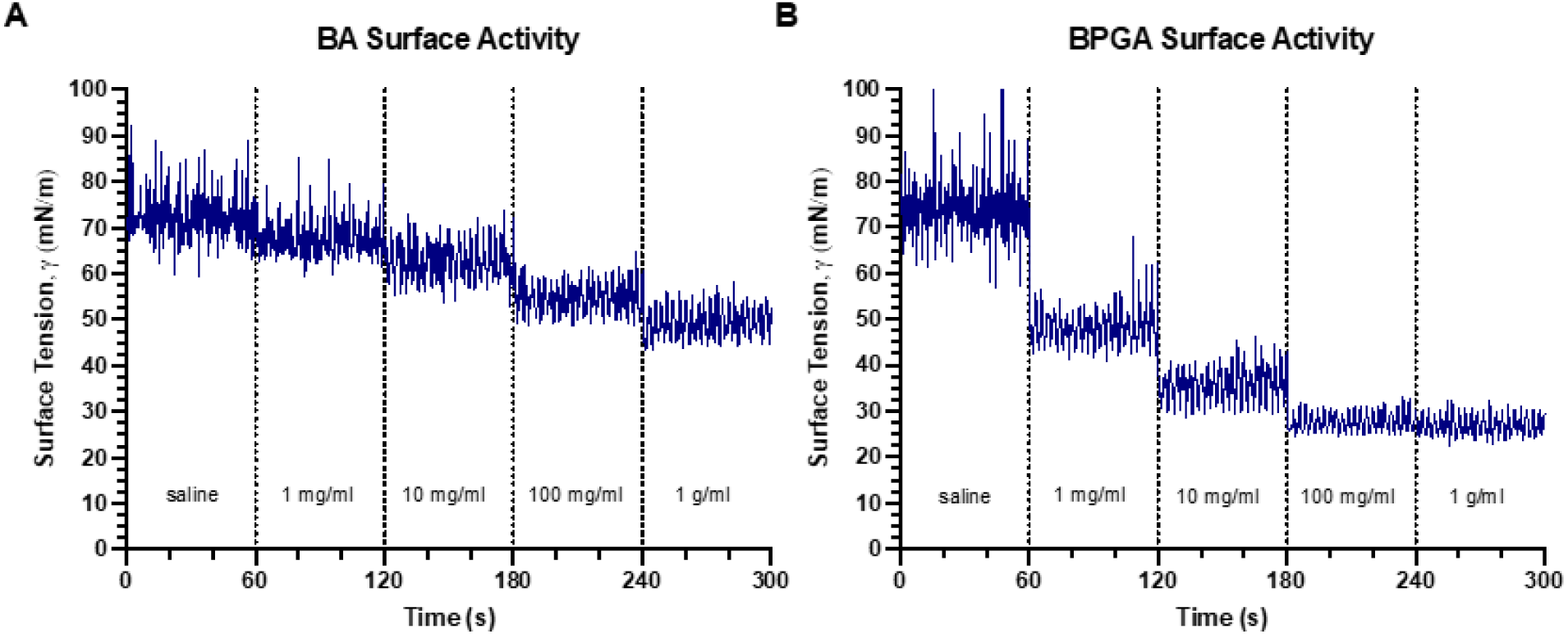
Vaping chemical surface activity. A) BA surface activity. B) BPGA surface activity. For both chemicals, concentrations were added to the CSD in ascending order, 1*µL* at a time, after each undergoing compression-expansion cycles for 60 seconds.

**Figure S4:**
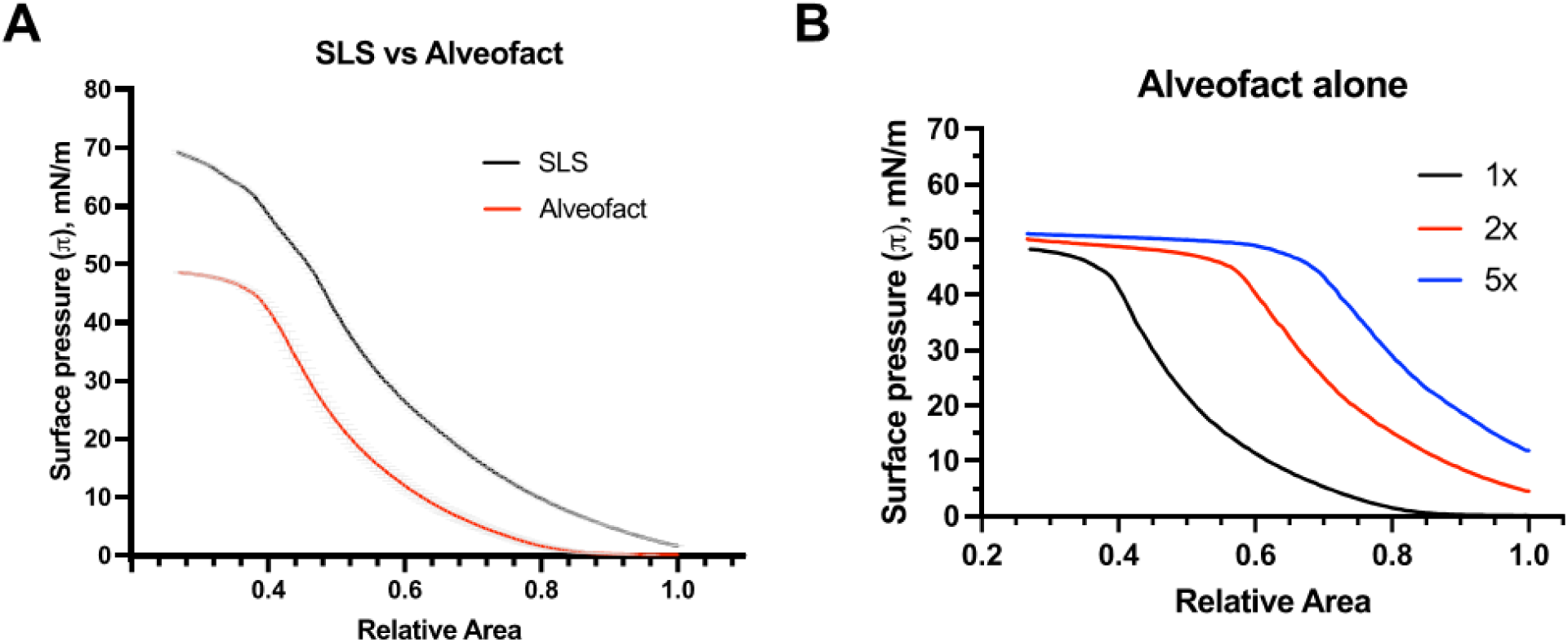
Clinical surfactant Alveofact *π − A* isotherms. A) SLS vs Alveofact on the LBT. *π − A* isotherms run with 20*µL* 1mg/ml of each surfactant at 150*cm*^2^*/min* on a 25°C saline subphase. Error bars represent standard error. N=3 replicates were produced for each condition. B) *π − A* isotherms for a range of Alveofact quantities. “1x” refers to 20*µL* 1mg/ml Alveofact. “2x” and “5x” refer to 2 times and 5 times molar quantity, respectively. N=3 for 1x, N=1 for 2x and 5x.

**Figure S5:**
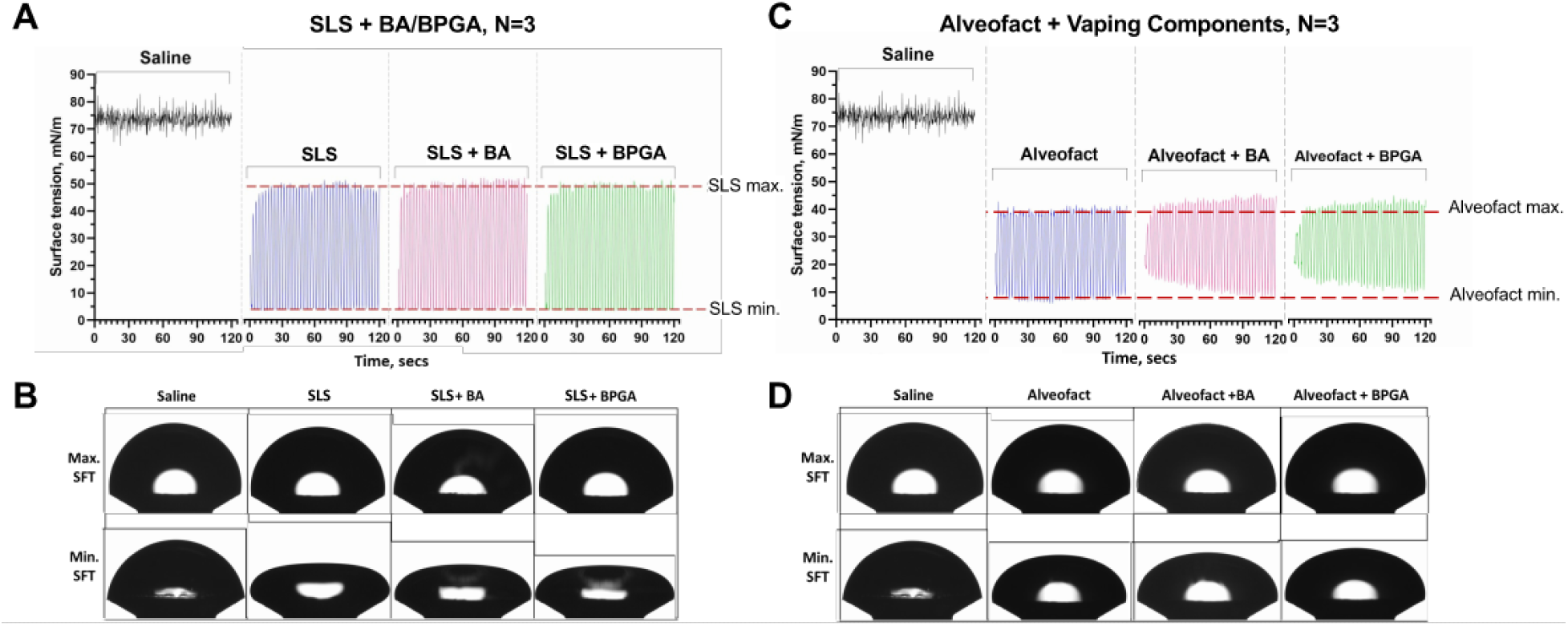
CSD surface tension-time cycles. A) SLS CSD compression-expansion cycles with BA and BPGA. Red lines outline the SLS alone maximums and minimums. B) Photographic representation of the Sessile Drop at maximum and minimum surface tension of the last cycle with conditions in A). C) Alveofact CSD compression-expansion cycles BA and BPGA. Red lines outline the Alveofact alone maximums and minimums. D) Photographic representation of the Sessile Drop at maximum and minimum surface tensions of the last cycle with conditions in C. The saline drop underwent cycles alone prior to the addition of any surfactant or vaping chemical. Cycles were completed at a rate of 20 cycles per minute over two minutes. N=3 replicates were collected for each condition.

**Figure S6:**
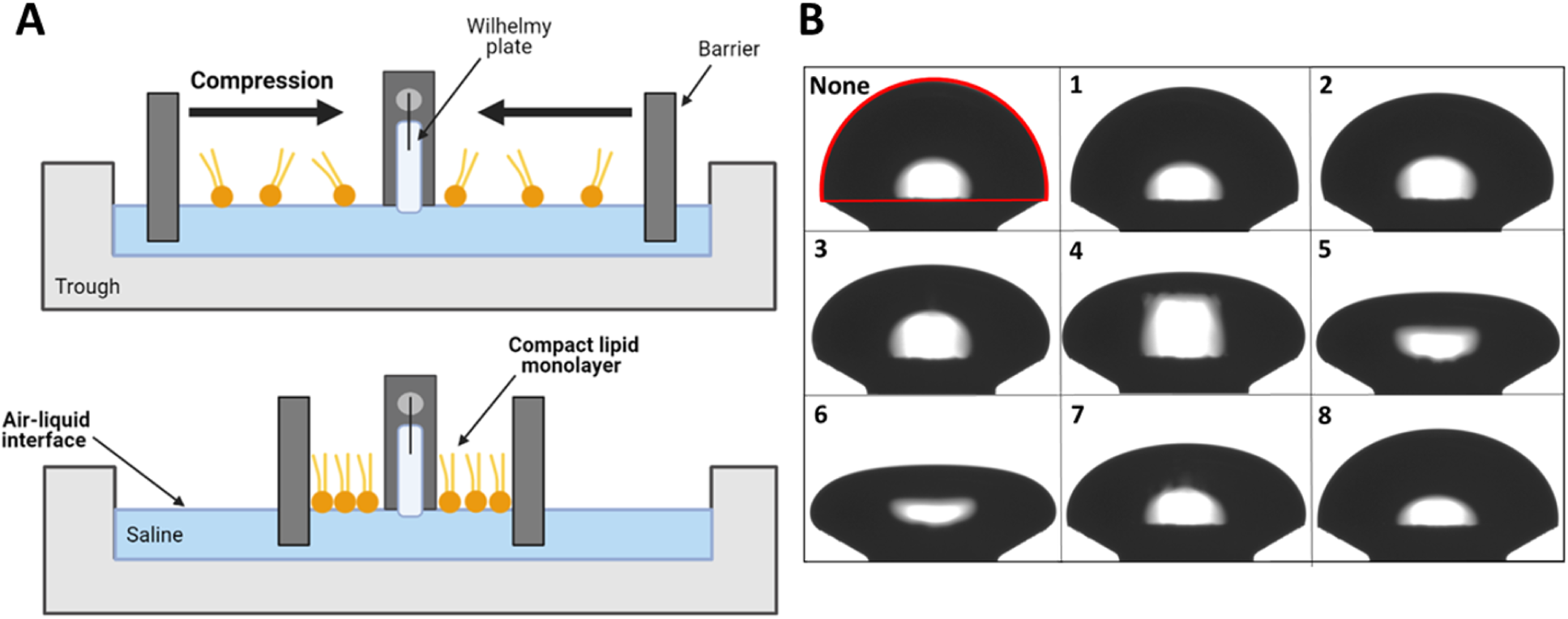
Overview of LBT and CSD techniques. A) A diagrammatic representation of the LBT set-up and *π − A* isotherm production. A surfactant film sitting at the air-liquid interface is compressed by reducing surface area via movement of motorised ribbon barriers to form a compact lipid monolayer. Surface pressure is recorded via the Wilhelmy plate surface pressure sensor and NIMA Software. B) Photographic representation of a CSD compression-expansion cycle. The drop, outlined in red, is sat on a stainless-steel pedestal (None). Surfactant can then be added and made to undergo compression-expansion cycles at physiological rates (1-8). The drop is compressed and expanded via the automated removal and replacement of subphase from beneath the pedestal. Surface tension is calculated from the contact angle between the pedestal and the drop surface in each image.

## TOC Graphic

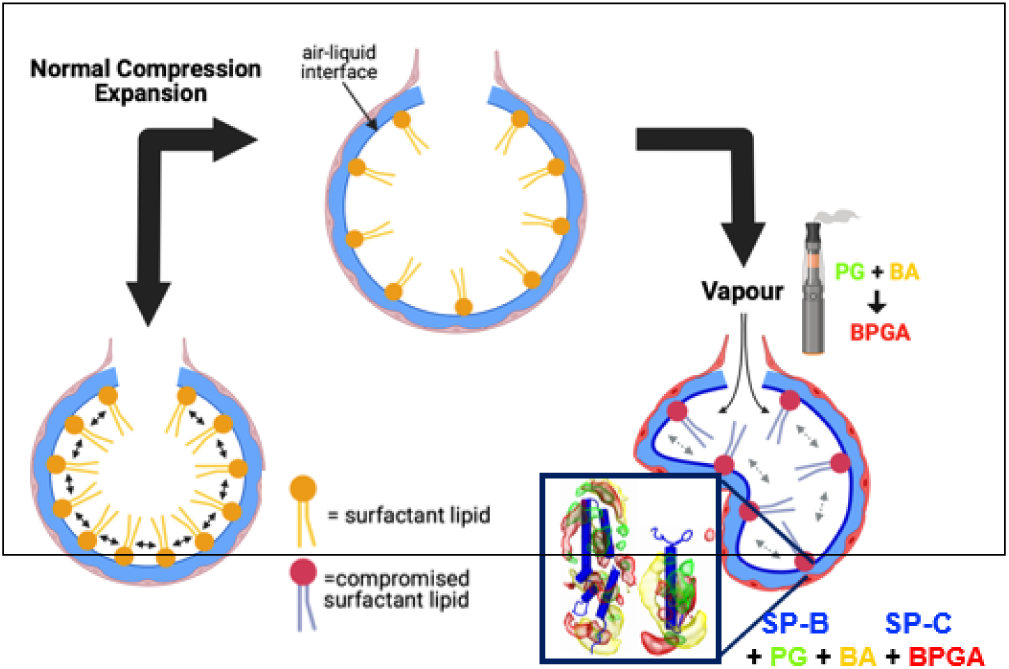

